# Identification of flavone and its derivatives as potential inhibitors of transcriptional regulator LasR of Pseudomonas *aeruginosa* using virtual screening

**DOI:** 10.1101/523381

**Authors:** Narek Abelyan, Hovakim Grabski, Susanna Tiratsuyan

## Abstract

Antibiotic resistance is a global problem nowadays and in 2017 the World Health Organization published the list of bacteria for which treatment are urgently needed, where *Pseudomonas aeruginosa* is of critical priority. Current therapies lack efficacy because this organism creates biofilms conferring increased resistance to antibiotics and host immune responses. The strategy is to “not kill, but disarm” the pathogen and resistance will be developed slowly. It has been shown that LasI/LasR system is the main component of the quorum sensing system in *P. aeruginosa*. LasR is activated by the interaction with its native autoinducer. A lot flavones and their derivatives are used as antibacterial drug compounds. The purpose is to search compounds that will inhibit LasR. This leads to the inhibition of the synthesis of virulence factors thus the bacteria will be vulnerable and not virulent. We performed virtual screening using multiple docking programs for obtaining consensus predictions. The results of virtual screening suggest benzamides which are synthetical derivatives of flavones as potential inhibitors of transcriptional regulator LasR. These are consistent with recently published experimental data, which demonstrate the high antibacterial activity of benzamides. The compounds interact with the ligand binding domain of LasR with higher binding affinity than with DNA binding domain. Among the selected compounds, by conformational analysis, it was found that there are compounds that bind to the same amino acids of ligand binding domain as the native autoinducer. This could indicate the possibility of competitive interaction of these compounds. A number of compounds that bind to other conservative amino acids ligand binding domain have also been discovered, which will be of interest for further study. Selected compounds meet the criteria necessary for their consideration as drugs and can serve as a basis for conducting further *in vitro / in vivo* experiments. It could be used for the development of modern anti-infective therapy based on the quorum sensing system of *P. aeruginosa*.

**Highlights:**

- Virtual screening using multiple docking programs for consensus predictions.
- Virtual screening reveal benzamides as potential inhibitors of LasR.
- Selected compounds bind to the same amino acids of LBD as the native autoinducer.
- Selected compounds meet the criteria necessary for their consideration as drugs.

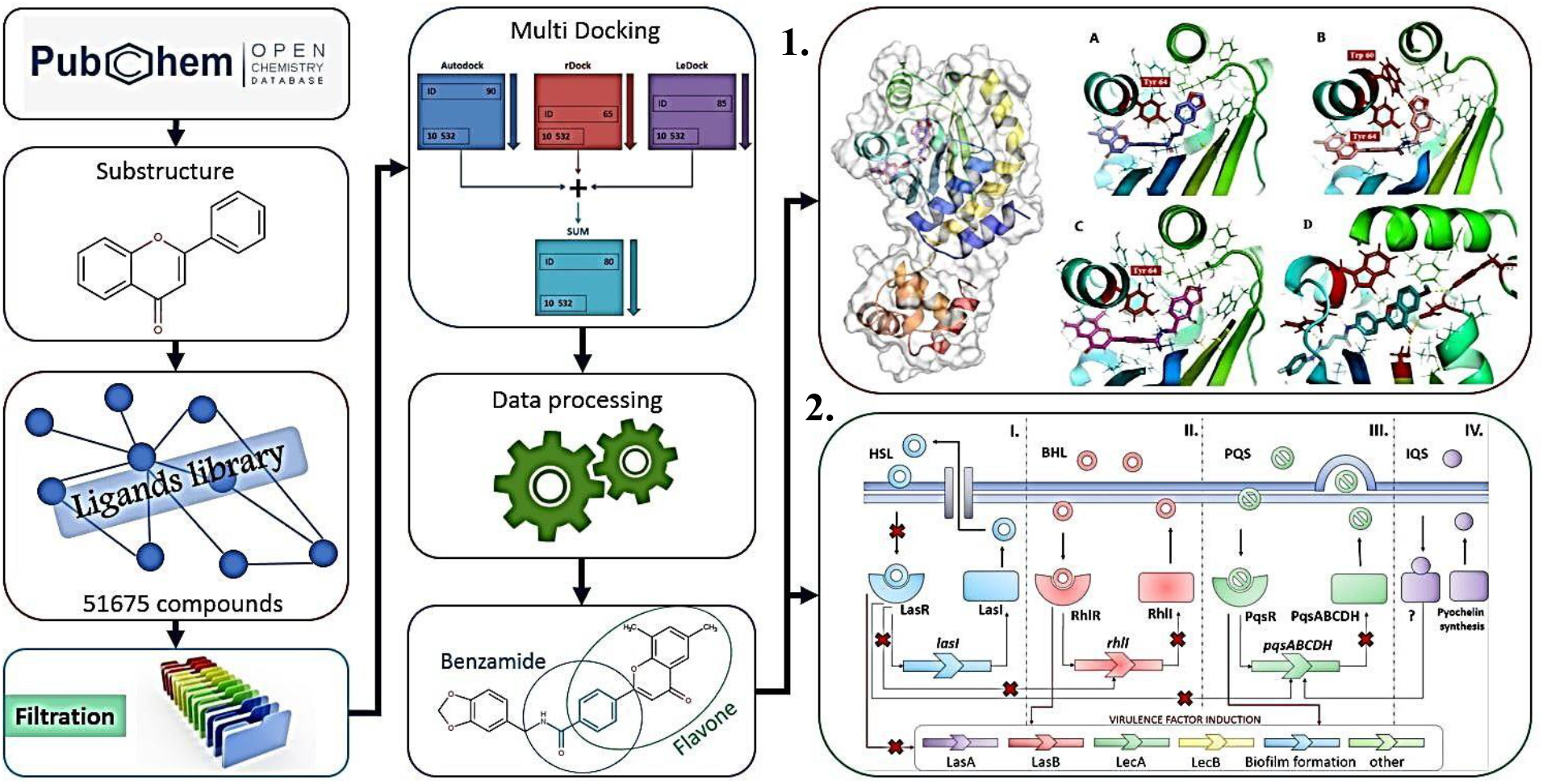
GRAPHICAL ABSTRACT. **1**: N- (1,3-benzodioxol-5-ylmethyl) -4- (6,8-dimethyl-4-oxochromen-2-yl) benzamide docking with LBD of LasR, A — ligand conformation predicted by AutoDock, B - by rDock and C - by LeDock,D - binding of CID 108754330 with LBD of LasR predicted by rDock. **2**: Four types of P. aeruginosa quorum sensing signaling systems I. Las, II. Rhl, III. Pqs and IV. IQS.

## 1. INTRODUCTION

Nowadays antibiotic resistance is a global problem, every year at least 2 million people get an antibiotic-resistant infection, and about 23,000 people die in the US [1]. In 2017, the World Health Organization (WHO) published the list of bacteria for which new antibiotics are urgently needed, where *Pseudomonas aeruginosa (P. aeruginosa)* is of critical priority [2]. These bacteria multiply in the host’s body, without any visible damage, but when they reach a certain concentration using quorum-sensing (QS) system, they unite into communities (biofilm). They secrete molecular signals, which are responsible for cellular communication, so they can jointly make decisions how to adapt to the environment and activate defense mechanisms. Biofilms promote the expression of virulence genes, including the genes responsible for antibiotics and multidrug resistance. Also, they represent a physical barrier that prevents the penetration of harmful substances such as antibiotics [3,4].

### 1.1. Mechanisms of P. aeruginosa quorum sensing

*P. aeruginosa* uses four types of quorum sensing signaling systems. Two of which are based on AHLs (acyl homoserine lactones): OdDHL (N- (3- oxododecanoyl) -L-homoserine lactone) and BHL (N-butanoyl-L-homoserine lactone). These autoinducers are recognized by two different signaling systems, and both of them includes a LuxR type receptor and a LuxI type synthase. OdDHL is synthesized by the synthase LasI and binds to the LasR receptor [5,6]. BHL is formed by RhlI synthase and is recognized by RhlR protein [7,8]. These two systems are integrated with the third Pseudomonas quinolone system (PQS), which uses quinolone as singnaling molecules [9]. The IV system is IQS (Integrated Quorum Sensing system). The IQS pathway remains unraveled and the IQS receptor is unknown. [10] Quorum sensing system of *P.aeruginosa* is hierarchical, where the activation of LasR leads to the stimulation of RhlR and PQS systems. Therefor LasR has a crucial role and is of greater interest for further research [11] (Fig. 1).

**Figure 1:**
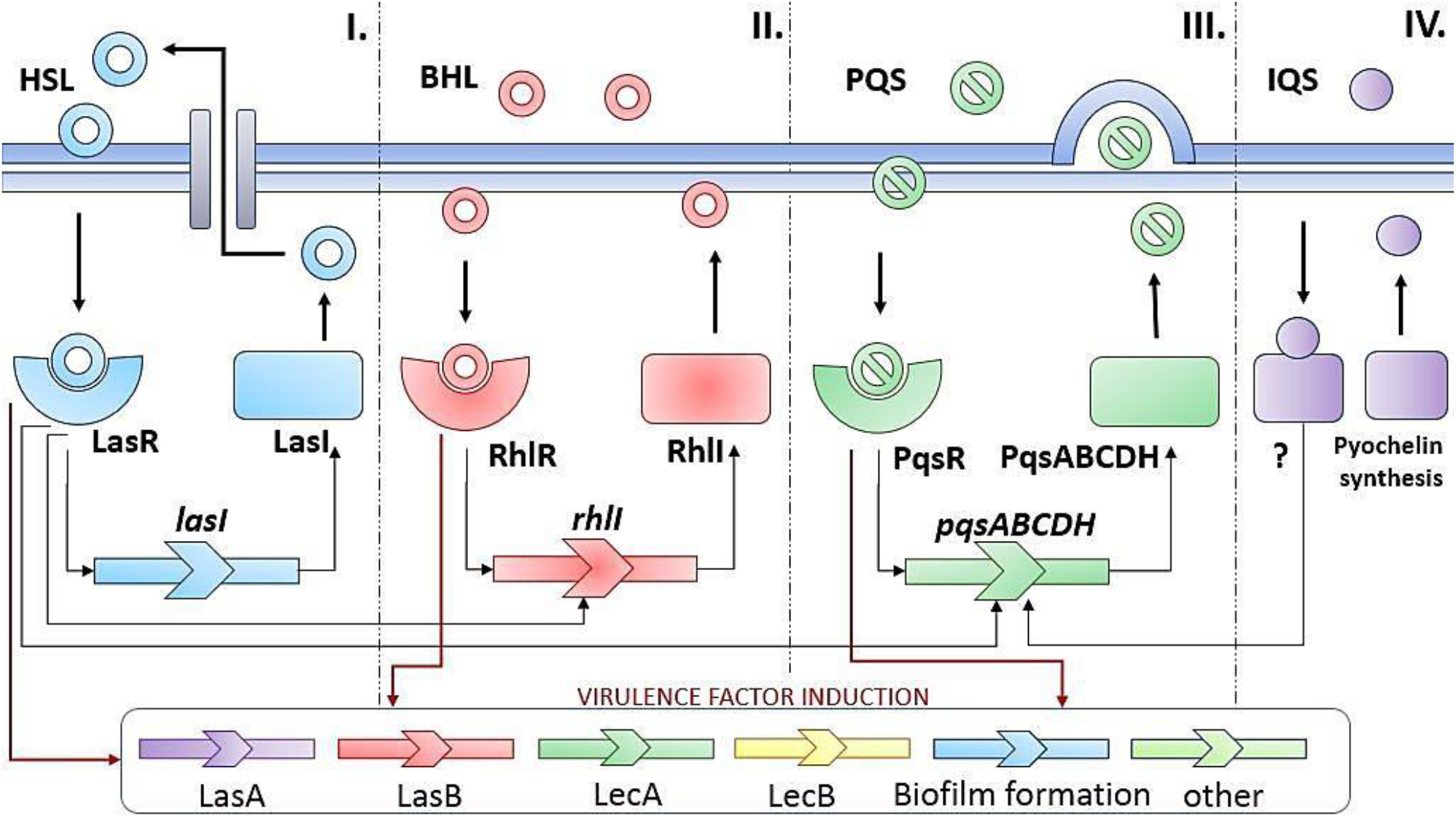
Mechanisms of P. aeruginosa quorum sensing: four types of quorum sensing signaling systems I. Las, II. Rhl, III. Pqs and IV. IQS.

### 1.2. Pathogen control strategy

In the presence of mature biofilms in *P. aeruginosa*, phagocytes fail to migrate to the extracellular polymer surface of biofilms. At the same time, bacteria can continue to ag- gressively react to the presence of leukocytes, regulating the production of a number of QS- controlled virulence factors, including rhamnolipids [12], pyocyanine [13], and exopolysac- charides [14].

The approach for improving this situation involves the development of compounds that do not kill the pathogenic bacteria by suppressing biosynthesis, but inhibiting or inactivating their virulence factors. In other words, the strategy is to “not kill, but disarm” the pathogen. It is assumed that resistance to such anti-virulent compounds will be developed more slowly, since they will not directly affect the viability of bacteria and will only affect their ability to infect humans.

The purpose of this work is to find compounds that will inhibit LasR functioning. This will lead to the suppression of biofilm formation and synthesis of virulence factors.

High throughput *in vitro* screening is one of the most effective and well-established methods used in modern drug design and discovery field. It is widely accepted to conduct preliminary use *in silico* screening, which reduces the number of test compounds for further *in vitro* and *in vivo* experiments, thereby reducing the high cost of the experiments. Also, it reduces the time of studies and increases the likelihood of detecting biological active compounds.

## 2. MATERIALS AND METHODS

### 2.1. Preparation of the protein structure and identification of binding sites

Due to the poor solubility of the LasR protein, crystallographic data of the complete structure of the receptor is not available. Therefore, the virtual reconstructed structure of the LasR receptor was used for the research [15]. The binding sites were also taken from the previous work [15] and verified by the MetaPocket server [16]. As a result, two binding pockets were identified: the ligand-binding domain (LBD) and the DNA-binding domain (DBD).

### 2.2. Preparation of the chemical compounds library

The PubChem - chemical compounds database [17], from which 51,675 flavones and their derivatives were selected using the substructure method, was used for the creation of low weight molecular ligands library. In addition library was filtered from compounds with poor adsorption and penetration, based on the Lipinski rule, which led to a decrease in the number of compounds to 21,232 [18]. Then Chemical Vendors filter was used to remove compounds without chemical manufacturers, which led to a reduction of the number of compounds to 10 532. For the preparation and the optimization of the ligands the conformational mobility (determination of the degrees of freedom) and charges for all atoms were calculated, polar hydrogen atoms were inserted. The structures were processed out using the software package OpenBabel v2.4.0 [19].

### 2.3. Molecular docking

Since the modern docking programs use simplified models of quantum chemical interac- tions, as well as lower resource and cost algorithms, for more reliable results, we used three docking programs — AutoDock Vina [20], rDock [21], and LeDock [22], which demonstrated high performance in a recent comprehensive assessment of docking programs [23].

At the initial stage, the binding affinities of the studied compounds with LasR protein with the two active sites were determined using the AutoDockVina software package [20]. AutoDockTools [24], was used for the configuration of the simulations. Using this program, two docking boxes were selected, whose sizes did not exceed 27,000 Å3, and “exhaustiveness” was set to 10, which is recommended for rapid virtual screening when using small boxes [25]. Table 1 shows the results of the docking analysis of the 10 compounds that showed the highest binding affinities in both sites.

**Table 1.** The results of the docking - analysis of compounds that showed the highest binding affinities with LBD and DBD of LasR

| **№** | **PubChem CID** | **Binding affinity-LBD (kcal/mol)** | **PubChem CID** | **Binding affinity -DBD (kcal/mol)** |
| --- | --- | --- | --- | --- |
| 1 | 108786995 | **-12.6** | 60617261 | **-10.9** |
| 2 | 108754324 | -12.3 | 46684079 | -10.5 |
| 3 | 108787038 | -12.2 | 108773117 | -10.2 |
| 4 | 108800047 | -12.2 | 16297521 | -10.2 |
| 5 | 1695677 | -12.0 | 24335408 | -10.1 |
| 6 | 108802999 | -12.0 | 16505442 | -10.0 |
| 7 | 108790351 | -11.9 | 108773119 | -10.0 |
| 8 | 108803145 | -11.9 | 31836827 | -10.0 |
| 9 | 108800021 | -11.8 | 41418450 | -10.0 |
| 10 | 108786994 | -11.8 | 52655891 | -9.9 |

It is important to note that the binding of the native ligand N-3-Oxo-Dodecanoyl-L-Homoserine lactone to LBD occurs with an energy of - 5.7 kcal. The results suggest that binding of selected compounds occurs intensively with both LBD and DBD. However, the majority of chemical compounds exhibit a higher affinity for LBD, so further research was carried out with this interaction site. To obtain more reliable results, molecular docking of the same 10,532 compounds was carried out by other docking programs as well. Molecular docking using rDock [21] includes three main steps: the first is system definition, the second is cavity generation, the third is docking. Fifty launches were performed for each ligand, as recommended by the developers of rDock [21]. To carry out molecular docking by the LeDock program [22], the input files were generated by the LePro program with default parameters [22].

Since three different docking programs were used, it became necessary to process a huge amount of molecular docking simulation data. For that purpose, we developed productive data processing approach.

### 2.4. Data processing

The first step involves the normalization of the obtained results and the creation of a common assessment system. Since different software packages calculate the interaction energy in different units, it became necessary to convert them into a single evaluation system, where 100 units receive a compound with the maximum amount of interaction energy, and 1 unit receives a compound with a minimum energy. For each program, the interaction energy of all compounds was converted to a common evaluation system. In Table 2 shows the normalized estimates of the LBD LasR binding of 10 compounds from the original list of 10 532 items.

**Table 2.** Normalized assessment of LBD LasR binding of compounds that showed high affinity for AutoDockVina [20], rDock [21] and LeDock [22].

| PubChem<br>CID | AutoDock | PubChem<br>CID | rDock | PubChem<br>CID | LeDock |
| --- | --- | --- | --- | --- | --- |
| 108786995 | 100 | 16801690 | 100 | 24280119 | 100 |
| 108754324 | 96.04 | 2817219 | 98.74 | 53486694 | 97.52 |
| 108787038 | 94.72 | 18568736 | 95.66 | 16408601 | 97.68 |
| 108800047 | 94.72 | 82046503 | 95.45 | 6336429 | 97.26 |
| 1695677 | 92.08 | 25555952 | 94.66 | 108809778 | 96.84 |
| 108802999 | 92.08 | 93713806 | 94.57 | 396728 | 96.84 |
| 108790351 | 90.76 | 21000649 | 94.51 | 16408064 | 96.63 |
| 108803145 | 90.76 | 18568755 | 94.48 | 108787045 | 95.58 |
| 108800021 | 89.44 | 93713804 | 93.28 | 5449792 | 95.37 |
| 108786994 | 89.44 | 108811376 | 92.98 | 16407858 | 94.95 |

The second step was the summation of the normalized estimates for each chemical compound obtained by the three programs. The third step was the ranking of compounds according to their total assessment, which includes compounds with a good predictive rating by the three programs (Table 3, Fig. S1). Thus, for those compounds that received false positive results with one program, the rank went down in the list. However, it is possible that all three programs will predict false positive results, but the probability of this is extremely small, since each program uses a specific algorithm for the prediction. In turn, the probability of a false prediction by each program become smaller compared to the probability of a true prediction. And since the purpose of virtual screening is to find a compromise between the share of truly active compounds in the top of the list and their total number in the sample, the use of this approach is quite reasonable.

**Table 3.** Compounds with good scores based on all the programs.

| IUPAC Name | PubChem | Autodock | rDock | LeDock | Sum |
| --- | --- | --- | --- | --- | --- |
| N-(1,3-benzodioxol-5-ylmethyl)-4-(6,8-dimethyl-4-oxochromen-2-yl)benzamide | 108786995 | 100 | 81.12 | 72.46 | 253.59 |
| N-[2-(3,4-dihydroxyphenyl)ethyl]-2-(5-hydroxy-4-oxo-2-phenyl-chromen-7-yl)oxyacetamide | 16408424 | 68.32 | 86.34 | 92.85 | 247.52 |
| N-(3-acetamidophenyl)-2-(5-hydroxy-4-oxo-2-phenylchromen-7-yl)oxyacetamide | 16407791 | 70.96 | 86.03 | 88.64 | 245.64 |
| 4-(6-methyl-4-oxochromen-2-yl)-N-[5-(2-methylpropyl)-1,3,4-thiadiazol-2-yl]benzamide | 108803142 | 85.48 | 76.67 | 82.34 | 244.50 |
| 4-(6-methyl-4-oxochromen-2-yl)-N-[2-(4-sulfamoylphenyl) ethyl]benzamide | 108790338 | 78.88 | 85.36 | 85.36 | 243.01 |
| 2-(5-hydroxy-4-oxo-2-phenylchromen-7-yl)oxy-N-(2-oxochromen-6-yl)acetamide | 16408830 | 65.68 | 83.23 | 92.01 | 240.92 |
| (2S)-2-[[2-(5-hydroxy-4-oxo-2-phenylchromen-7-yl) oxyacetyl ]amino]-3-(1H-indol-3-yl)propanoic acid | 29147486 | 73.6 | 78.45 | 88.22 | 240.28 |
| 3-[[[4-(6,8-dimethyl-4-oxochromen-2-yl)benzoyl] amino]methyl]benzoic acid | 108800059 | 88.12 | 71.76 | 79.61 | 239.49 |
| 2-(5-hydroxy-4-oxo-2-phenylchromen-7-yl)oxy-N-(pyridin-3-ylmethyl)acetamide | 16408601 | 67.0 | 74.34 | 97.68 | 239.03 |
| N'-(4-fluorophenyl)-N-[4-(6-methoxy-4-oxochromen-2-yl)phenyl]butanediamide | 108754330 | 81.52 | 85.79 | 70.57 | 237.89 |

A script were written for the automatization of the entire data processing and analysis. The scheme of the algorithm is shown in (Fig. 2).

**Figure 2:**
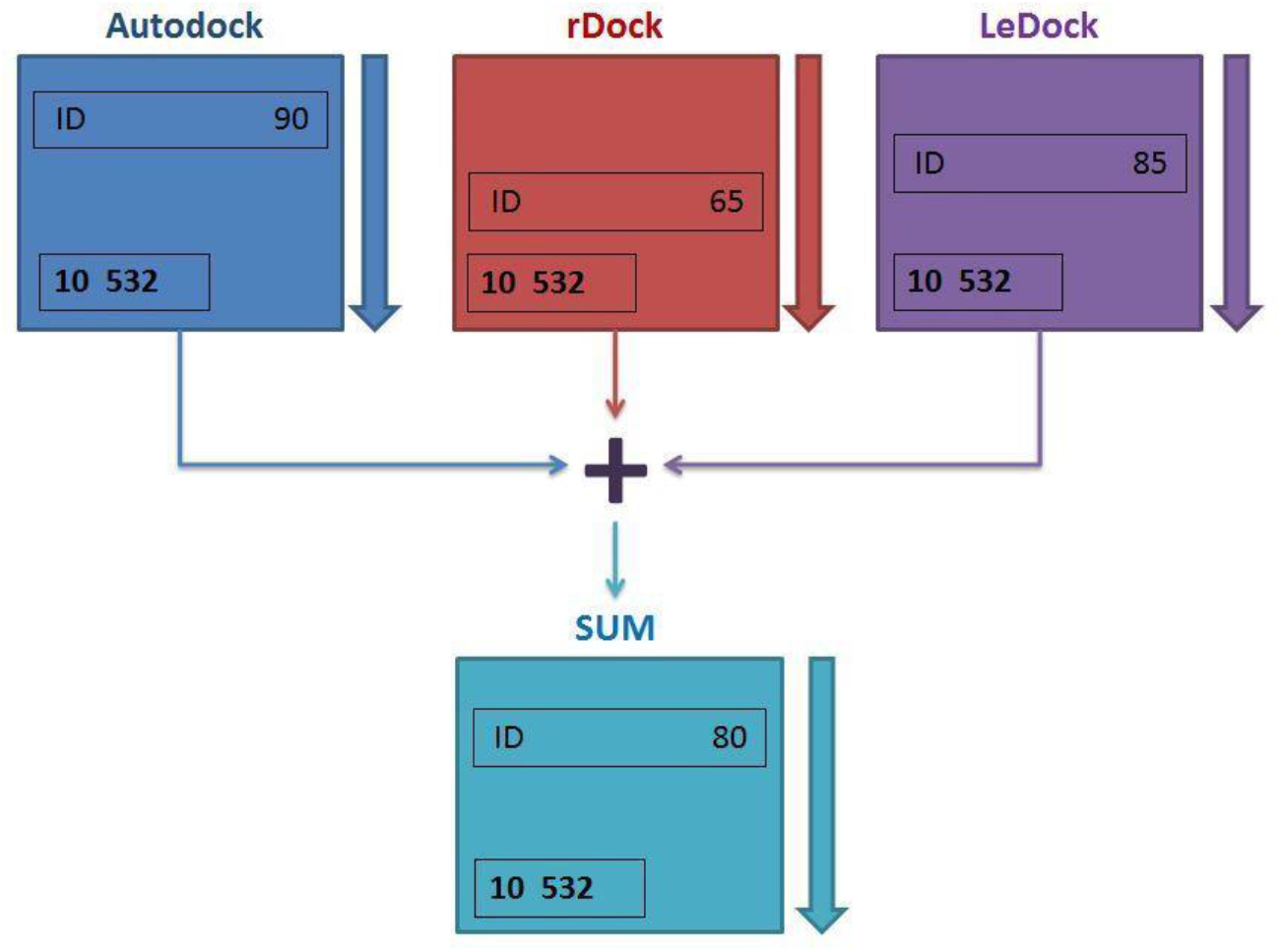
Scheme of the algorithm

### 2.5. Similarity analysis of drugs

Calculation of physicochemical descriptors, as well as the prediction of ADME properties: Absorption, Distribution, Metabolism, Excretion and Toxicity, were determined using three different programs: SwissADME [26], admetSAR [27] and PASS online [28], which are used for drug discovery. The toxicity prediction was also carried out using PASS online [28], which estimates the predicted types of activity with the estimated probability for each type of activity “to be active” Pa and “to be inactive” Pi, which vary from zero to one [28].

## RESULTS AND DISCUSSION

The 10 compounds that received high marks for the interaction with LBD LasR, based on the results of three programs, were selected from 10532 tested compounds (see Fig. S1 in additional materials on the journal’s website - molecbio.com).

- The structure of the native ligand (HLS) has an amide bond [-CO-NH-], which is responsible for the interaction with Asp73 and Tyr56 LBD LasR. The same chemical bond is present in some of the compounds that are known to interact with LasR [34–37]. It should be noted that in all of the 10 compounds identified by us there was an amide bond. This may indicate comparability of the obtained results and the importance of this bond that ensures strong interaction with LBD LasR. The leading compounds identified by us also have structural similarities with compounds for which the inhibitory effect of the lasR / lasI system has already been identified.
- 7 of the 10 compounds contain a benzamide group. In three of them, the benzene ring is linked to the nitrogen atom of the amino group (CID 16407791, CID 16408830, CID 108754330) and in four of the compounds the benzene ring is linked to the carbon of the carbonyl group (CID 108786995, CID 108803142, CID 108790338, CID 108800059). From the benzamides, CID: 108786995 received the maximum rating based on the results of three programs (Table 3, Fig. 3). There are experimental data that shows that compounds a benzamide group have anti-virulent properties both in vitro and in vivo models. In the work of Yang et al. [38], a compound containing a benzamide group, Nifuroxazide, exhibited a QS inhibitory effect, with a direct inhibition of LasR. In the work of Ute et al. [37], Triphenyl mimics of 3OC12-HSL were found, which interact with LasR and they also contain benzamide group. In the work of Melissa et al. [36], as a result of screening, a number of compounds with benzamide group that have anti-virulent properties were identified. These compounds reduced the production of HAQ, which is synthesized by the PqsR system. The most potent inhibitors (M51, M34, M62, M50, and M64) did not inhibit proteins involved in the direct synthesis of HAQ (expressed by the pqsABCD genes), thus suggesting that these inhibitors target MvfR or another upstream regulatory component. The interaction of M64 with MvfR has been shown, which may lead to a decrease in HAQ production. It should be noted that the decrease in HAQ production and the activation of the cascade of the PqsR system (MvfR) also occurs through the LasI / LasR system. And since the majority of identified potential inhibitor candidates contain a benzamide group, it can be assumed for the rest of the compounds (M51, M34, M62, M50) that the decrease of HAQ production was due to the inhibition of the LasR / LasI system. According to the article by Yang et al. [38], the 3 identified antiviral compounds, including Nifuroxazide, besides the direct inhibition of LasR, also had a direct inhibitory effect on the Rhl and PqsR quorum-sensing systems. Comparing the existing literature data, it can be assumed that benzamide-containing compounds have the potential to inhibit various pathways of the regulatory network of the QS system. Therefore, the compounds identified by us in the future can potentially inhibit simultaneously two QS systems.
- CID29147486 contains benzene ring associated with pyrrole (indole). The same chemical structure is found in indol-AHL. It is a compound that inhibits LasR protein [39].
- CID 108800059 contains the structure of benzoic acid, which is part of salicylic acid, which also has an inhibitory effect on LasR [38].
- CID 16408830 contains the coumarin group (Coumarin (1,2-Benzopyrone)). It has been shown that coumarin has anti-biofilm activity against P. aeruginosa [40].

**Figure 3.**
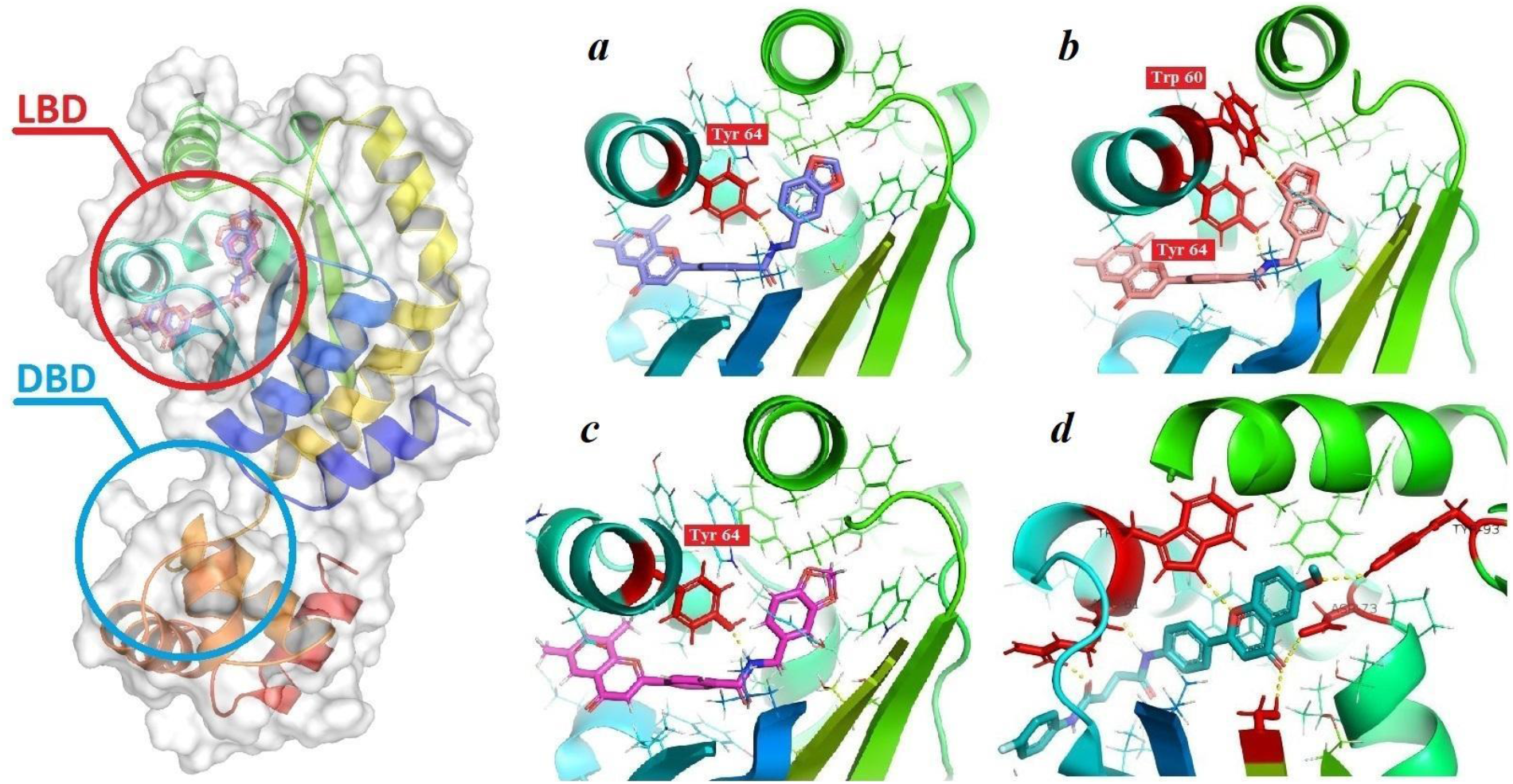
CID 108786995 docking with LBD of LasR, *a* − ligand conformation predicted by AutoDock, *b* − by rDock and *c* − by LeDock, *d* − binding of CID 108754330 with LBD of LasR predicted by rDock. For the visualization of the results, MarvinSketch [41], PyMOL[42] and VMD [43] software packages were used.

### Conformational analysis

Hydrogen and hydrophobic interactions analysis and visualization were performed using LigPlot+ [44] program for the 10 hit compounds from the ranked list. Among the selected compounds, by conformational analysis, it was found that there are compounds that bind to the same LBD amino acids as the native ligand. It is known from crystallographic data that N-3-Oxo-Dodecanoyl-L-Homoserine lactone forms hydrogen bonds with Asp73, Trp60, Tyr56, Ser129 [34] in the LBD pocket (Fig. 4). This could indicate the possibility of competitive interaction of these compounds. It should be also noted that ligands were also found that can strongly bind with other conservative amino acids due to the formation of hydrogen bonds or van der Waals forces and electrostatic interaction (see Fig. S2-S4 in additional materials on the journal’s website - molecbio.com).

**Figure 4.**
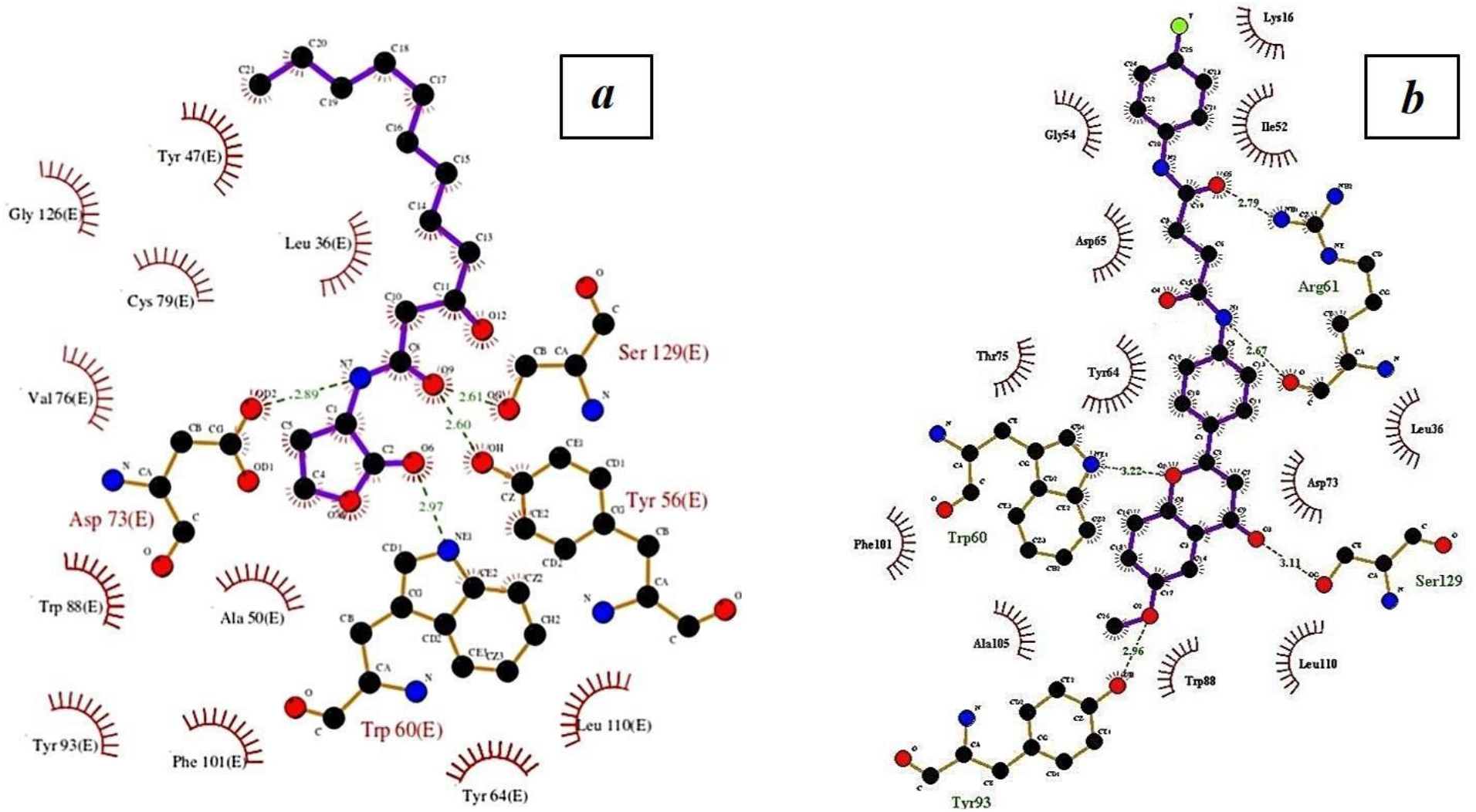
*a* − The crystallographic structure of the native ligand N-3-Oxo-Dodecanoyl-L-homoserine lactone with LBD of LasR; *b* − binding of CID 108754330 with LBD of LasR predicted by rDock.

The analysis of the conformations obtained using AutoDock Vina [23] demonstrate that compounds CID 29147486, 16408424, 16407791, 108790338, 108803142, 108786995 and 108800059 form hydrogen bonds with LBD of LasR. From the mentioned compounds only CID 108790338, 108803142, 108786995, 108800059 form a hydrogen bond with the conserved amino acid residue Tyr64 of LBD LasR, while CID 29147486 forms a hydrogen with the semi conserved Ala50 (Table 4).

**Table 4.**
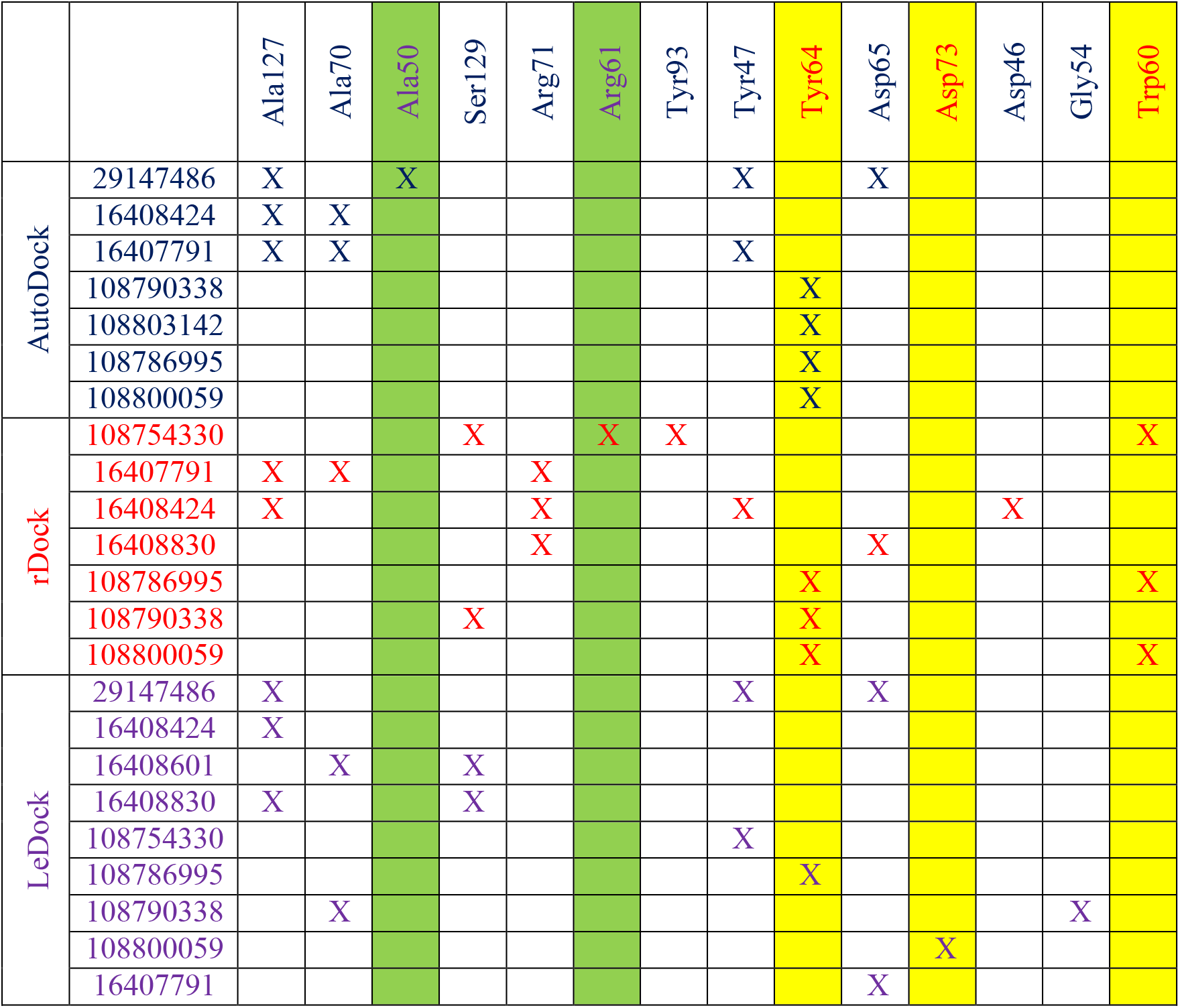
The interaction of ligands with amino acids of the LBD binding site. Yellow columns correspond to highly conserved amino acids, green - to semi-conservative amino acids.

Analysis of the conformations obtained using rDock [24] simulations indicated that CID 16407791, 16408424, 16408830, 108754330, 108786995, 108790338, 108800059 form hydrogen bonds with the LBD of the transcriptional regulator. Compounds CID 108754330, 108786995, 108800059 form a hydrogen bonds with conserved amino acid residue Trp60 of LBD of LasR, while CID 108786995, 108790338, 108800059 with Tyr64 and CID 108754330 with Arg61.

The results of the analysis of the conformations from LeDock [25] simulations demonstrate that CID 29147486, 16408424, 16408601, 16408830, 108754330, 108786995, 108790338, 108800059, 16407791 form hydrogen bonds with LBD of LasR. From the mentioned compounds CID 108800059 forms hydrogen bonds with conserved Asp73, while CID 108786995 forms with Try64 with LBD of LasR.

### Similarity analysis of drugs

The parameters of the CID 108786995 compound indicate that its potential properties can be used as a drug in biological systems (Fig. S5). The admetSAR package [32], predicted that this compound has high probability of adsorption through the gastrointestinal tract (HIA+) and the hemato-encephalic barrier (BBB+). SwissADME [31] also predicted the ability to adsorb through the gastrointestinal tract (HIA+), but it does not indicate that it can pass through the hemato-encephalic barrier (BBB-). The toxicity prediction was carried out using PASS online [33]. The predicted results of CID 108786995 show that it is not toxic. Selected compounds satisfy the criteria for drugs similarity and additional *in vitro* and *in vivo* experiments should be performed.

## CONCLUSIONS

Most flavones and their derivatives can be considered as potential antibacterial drug compounds, which interact with the LBD domain with higher binding affinity than with DBD. Among the selected compounds,based on conformational analysis, it was found that there are compounds that bind to the same amino acids from LBD as the native ligand. This could indicate the possibility of competitive interaction of these compounds. A number of compounds that bind to other conserved amino acids from LBD have also been discovered, which will be of interest for further study. The results of virtual screening are consistent with experimental data, which demonstrate the high antibacterial activity of benzamides [35–37] Comparing the existing literature data, it can be assumed that benzamide-containing compounds have the potential to inhibit various pathways of the regulatory network of the QS system. Therefore, the compounds identified by us in the future can potentially inhibit simultaneously two QS systems. Selected compounds meet the criteria necessary for their consideration as drugs and can serve as a basis for conducting further *in vitro / in vivo* experiments. It could be used for the development of modern anti-infective therapy based on the QS system of *P. aeruginosa*.

## Acknowledgments

The research was done within the Ministry of Education and Science of the Republic of Armenia, State Committee of Science: 10-2-1-4, Government budget financing.

