## Supplementary material for "Identification of flavone and its derivatives as potential inhibitors of transcriptional regulator LasR of Pseudomonas *aeruginosa* using virtual screening"

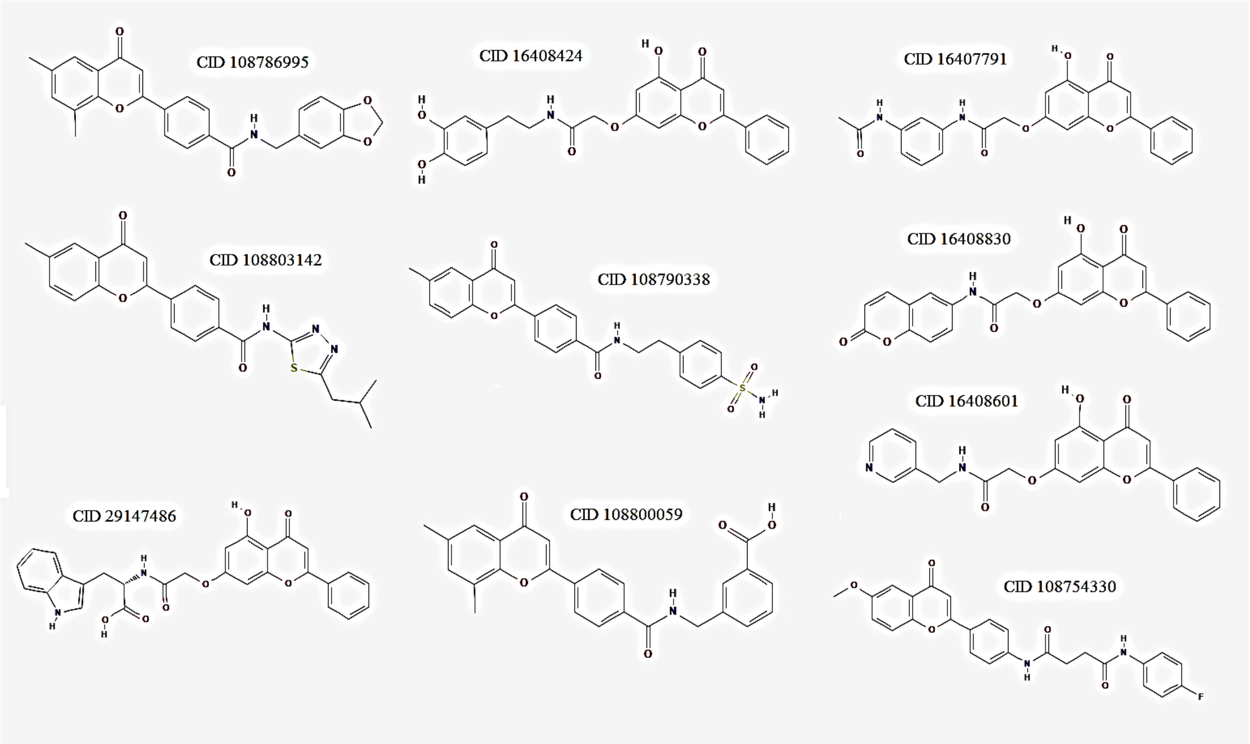


*Fig. S1: 10 compounds that received high scores based on the three programs results*

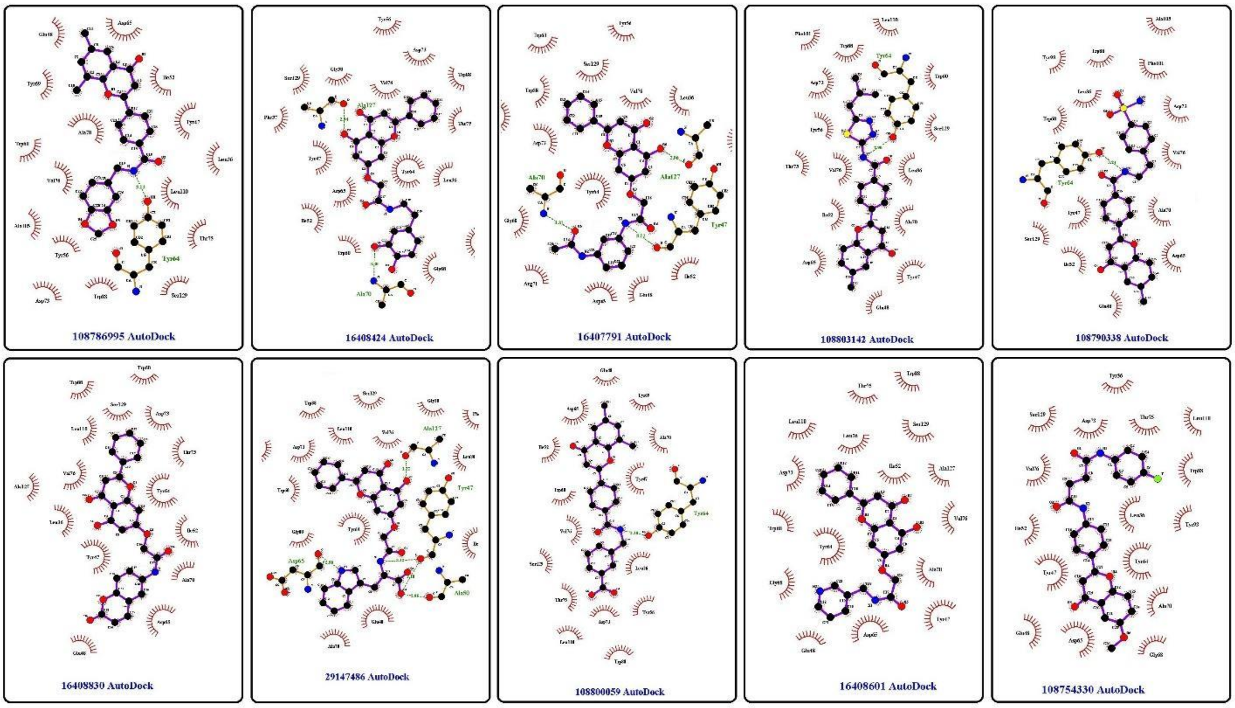


*Fig. S2: Hydrogen and hydrophobic interactions of the selected compounds with LBD of LasR obtained using Autodock Vina*

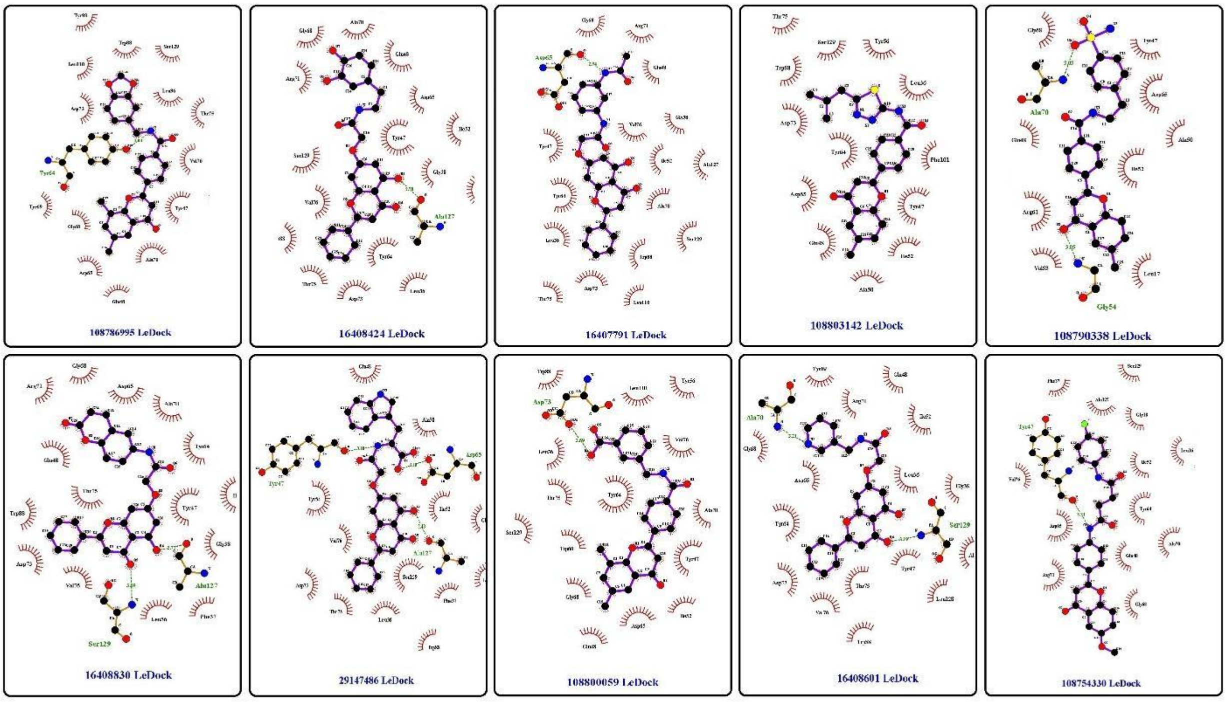


*Fig. S3: Hydrogen and hydrophobic interactions of the selected compounds with LBD of LasR obtained using LeDock*

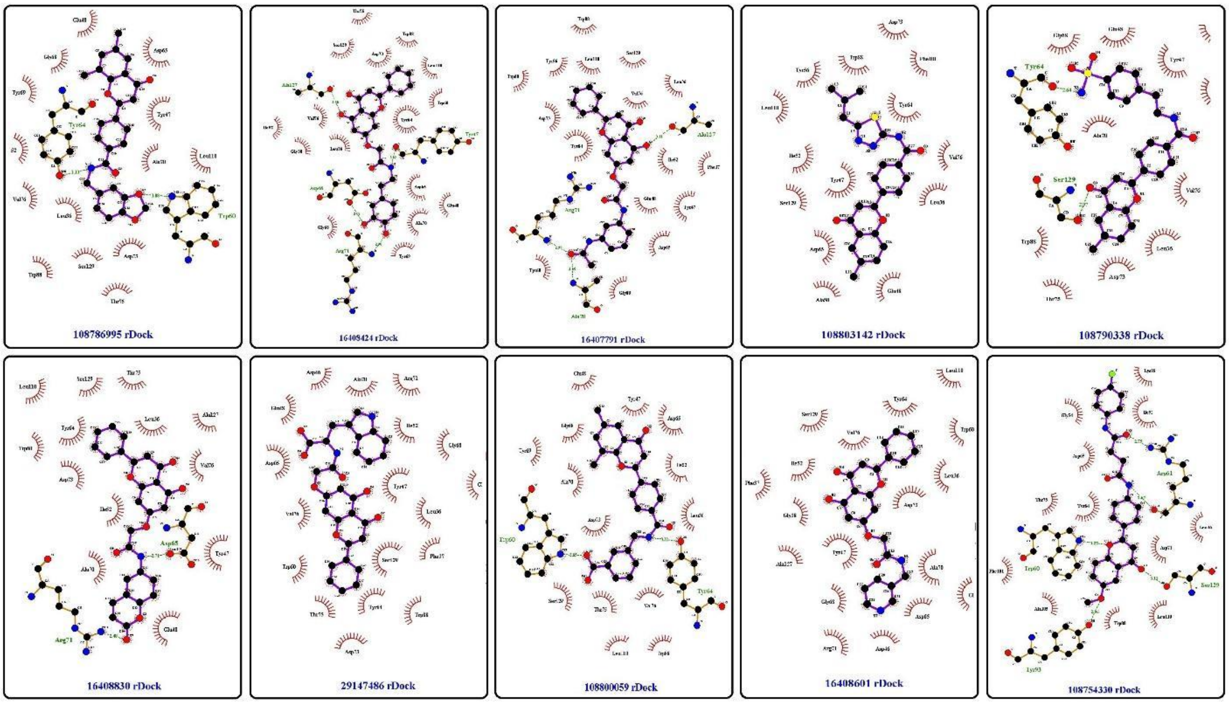


*Fig. S4: Hydrogen and hydrophobic interactions of the selected compounds with LBD of LasR obtained using rDock*

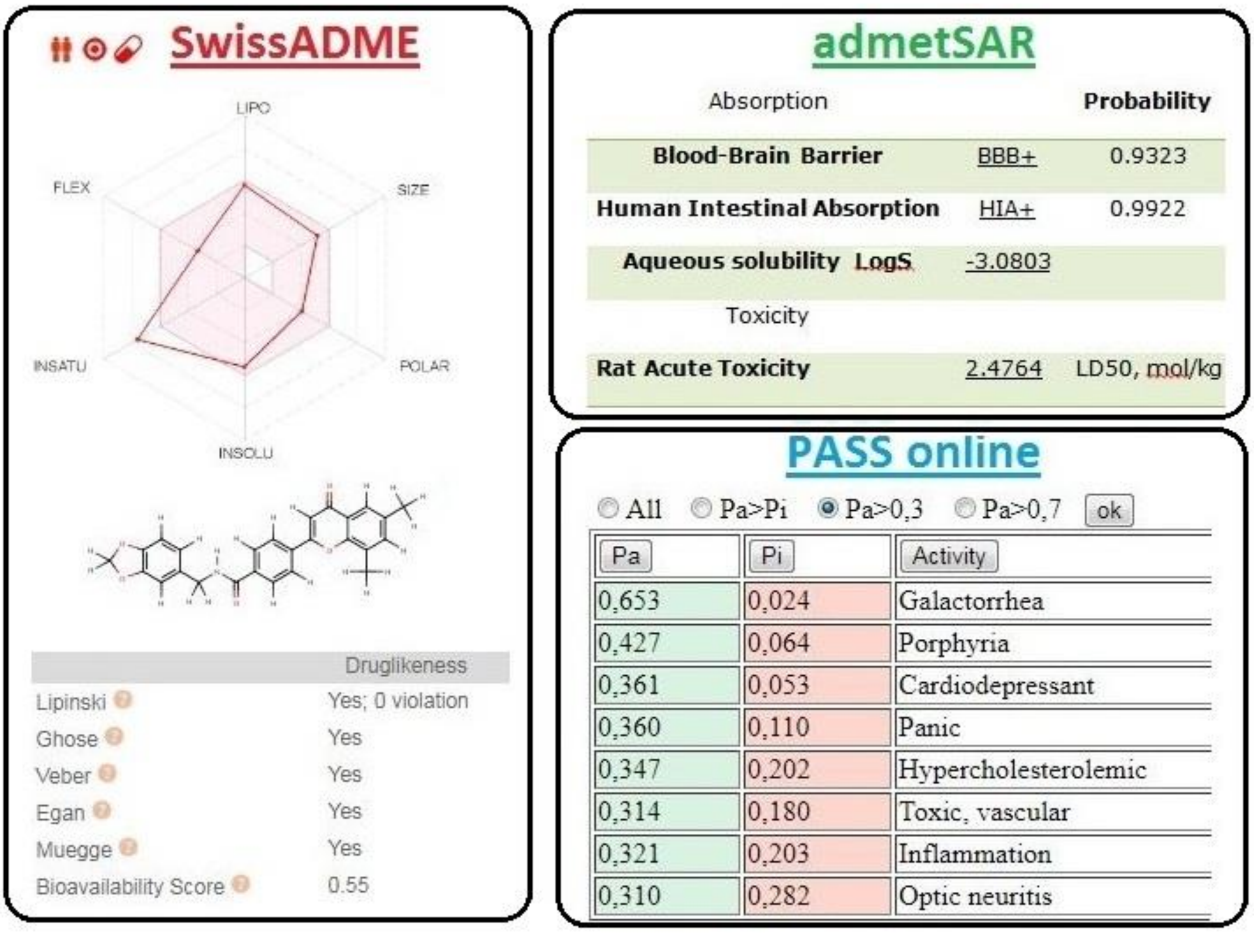


*Fig. S5: Similarity analysis of drugs*
